# Multi-Step Control of Homologous Recombination by Mec1/ATR Ensures Robust Suppression of Gross Chromosomal Rearrangements

**DOI:** 10.1101/2023.11.21.568146

**Authors:** Bokun Xie, Ethan James Sanford, Shih-Hsun Hung, Mateusz Maciej Wagner, Wolf-Dietrich Heyer, Marcus B. Smolka

## Abstract

The Mec1/ATR kinase is crucial for genome stability, yet the mechanism by which it prevents gross chromosomal rearrangements (GCRs) remains unknown. Here we find that in cells with deficient Mec1 signaling, GCRs accumulate due to the deregulation of multiple steps in homologous recombination (HR). Mec1 primarily suppresses GCRs through its role in activating the canonical checkpoint kinase Rad53, which ensures the proper control of DNA end resection. Upon loss of Rad53 signaling and resection control, Mec1 becomes hyperactivated and triggers a salvage pathway in which the Sgs1 helicase is recruited to sites of DNA lesions via the 911-Dpb11 scaffolds to favor heteroduplex rejection and limit HR-driven GCR accumulation. Fusing an ssDNA recognition domain to Sgs1 bypasses the requirement of Mec1 signaling for GCR suppression and nearly eliminates D-loop formation, thus preventing non-allelic recombination events. We propose that Mec1 regulates multiple steps of HR to prevent GCRs while ensuring balanced HR usage when needed for promoting tolerance to replication stress.

## Introduction

Gross chromosomal rearrangements (GCRs) are aberrant structural variations in chromosomes, such as deletions, translocations and amplifications that compromise genomic stability and drive oncogenesis ^1–3^. An important source of GCRs is DNA replication stress. The progression of replication forks is often impeded by various types of barriers such as DNA lesions, difficult-to-replicate regions and transcriptional intermediates, leading to stalled replication fork structures that can be converted into double-strand breaks (DSBs) through the action of nucleases ^4–9^. During DNA replication, DSBs are commonly repaired via homologous recombination (HR), a multi-step process that includes DNA end resection, strand invasion, DNA synthesis and the processing of recombination intermediates ^10–12^. HR is a high-fidelity mode of DNA repair, helping to prevent genomic rearrangement and maintain the overall integrity of the genome when sister chromatids are used as templates. However, when strand invasion occurs at the wrong locus, non-allelic HR between partially homologous (homeologous) sequences can happen, leading to the formation of GCRs ^13,14^. This includes formation of heteroduplex DNA between the homeologous sequences which is subject to recognition by the mismatch repair system and heteroduplex rejection to avert GCRs ^15^. How cells regulate HR-mediated DNA repair to prevent non-allelic recombination and GCRs is not fully understood. In particular, how cells balance the use of HR to ensure its adequate use while discerning from contexts where it may drive GCR events is a complex problem that likely requires decision-making steps and sensing mechanisms.

Understanding the mechanisms of GCR suppression in higher eukaryotes is challenged by the lack of sensitive and effective assays for monitoring GCRs that can be coupled to genetic screens. In contrast, significant progress in the study of the genesis of GCRs has been made using the “classical” GCR assay based on canavanine and 5-fluoroorotic acid (5-FOA) selection in *S. cerevisiae* to screen for spontaneous GCRs associated with the combined loss of the *CAN1* and *URA3* genes placed at the non-essential left arm of chromosome V ^16^. Using this approach, numerous factors implicated in DNA repair and DNA damage checkpoint have been identified to play pivotal roles in suppressing GCRs, among them the Mec1/ATR and the Tel1/ATM kinases ^13,17–19^. While deletion of *MEC1* leads to significant increases in GCR rates (∼200 fold higher compared to WT) and deletion of *TEL1* has no effects on GCR rates, *mec1*Δ *tel1*Δ cells display one of the highest GCR rates reported (over 10,000 fold increase compared to WT) ^19^. Despite the crucial roles of Mec1 and Tel1 in GCR suppression, the mechanism by which these kinases prevent GCR accumulation remains incompletely understood.

Mec1 is a phosphoinositol-3-Kinase-like kinase (PIKK) that functions as a sensor of DNA replication stress by recognizing single-strand DNA (ssDNA) accumulation mainly at stalled replication forks and recessed DSBs ^20–23^. Mec1 recognizes replication protein A (RPA)-coated ssDNA via its cofactor Ddc2 ^20,24^ and, once recruited, is activated by proteins such as Dpb11, Ddc1 and Dna2 that contain a disordered Mec1-activating domain ^25–28^. Active Mec1 phosphorylates and activates the downstream kinase Rad53 to initiate the canonical DNA damage checkpoint response that promotes cell cycle arrest, fork stabilization and protection, inhibition of origin firing, regulation of dNTP production and transcriptional reprogramming ^29–33^. The classical checkpoint adaptor Rad9 contributes to transducing signaling from Mec1 to Rad53, while also playing roles in the control of DNA end resection, the first step in HR-mediated DNA repair ^34–36^. Mec1 has also been reported to play roles in the regulation of HR-mediated DNA repair independently of its canonical function in checkpoint signaling ^37–41^. Depending on the context, Mec1 can exert inhibitory or stimulatory effects on DNA end resection control. For example, while early in the response to DNA lesions Mec1 can inhibit resection by facilitating the recruitment and oligomerization of the resection antagonist Rad9 at DNA lesions ^36,42^, at later stages Mec1 can then promote long-range resection by mediating the recruitment of the DNA repair scaffolding protein Slx4, which counteracts the resection block formed by Rad9, therefore promoting resection ^43,44^. The recruitment of both Rad9 and Slx4 relies on their interaction with Dpb11, a multi-BRCT domain scaffold that recognizes phosphorylated Rad9 or Slx4 and stabilizes them at DNA lesions ^45,46^. In addition to resection control, Mec1 regulates strand exchange through the phosphorylation of the strand exchange factor Rad55 ^47,48^ and of the recombinase Rad51 ^38^. Mec1 phosphorylation has been proposed to control the ATPase activity of Rad51 and influence HR ^38^.

The ability of Mec1 to suppress GCRs is largely independent of its canonical function in activating the DNA damage checkpoint ^19,49^. This is best evidenced by the lower rates of GCRs in cells lacking *RAD53* compared to the rates observed in cells lacking *MEC1* ^19^. Despite strong genetic evidence pointing to a crucial checkpoint-independent role for Mec1 in GCR suppression, the precise mechanism by which Mec1 signaling promotes such suppression remains unknown.

To characterize the checkpoint-independent role of Mec1 in GCR suppression, here we monitored Mec1 signaling in *rad53*Δ cells using phosphoproteomics and find that loss of the DNA damage checkpoint triggers hyper-activation of Mec1 signaling and hyper-phosphorylation of the Sgs1 helicase, a helicase involved in multiple steps of HR, including resection, heteroduplex rejection and dissolution ^50–53^. In checkpoint defective cells, GCRs are largely suppressed by Mec1-dependent recruitment of Sgs1 to sites of DNA lesions via phosphorylation of the 9-1-1 clamp and Sgs1, which assembles a 911-Dpb11-Sgs1 complex that increases heteroduplex rejection. Fusing an ssDNA recognition domain to Sgs1 (RBD-Sgs1 chimera) bypasses the requirement of Mec1 signaling for GCR suppression and nearly eliminates D-loop formation, consistent with a model that Mec1 suppresses GCRs by promoting heteroduplex rejection and HR quality control, thus preventing non-allelic recombination events. We propose that Mec1 prevents GCRs through a redundant system of HR control involving both resection control via checkpoint activation and heteroduplex rejection via Sgs1 recruitment and regulation. GCRs drastically rise in cells lacking *MEC1* due to the abolishment of both GCR suppressing functions.

## Results

### Loss of *RAD53* or *RAD9* triggers Mec1 hyper-activation and dependency on Sgs1 for GCR suppression

Loss of MEC1 causes a ∼200 fold increase in the rates of GCRs, while cells lacking RAD53 exhibit only a ∼30 fold increase in GCR rates ^19^. Since the loss of RAD53 impairs fork stabilization and resection control, which are expected to contribute to promoting GCR events, we reasoned that Mec1 must promote GCR suppression in rad53Δ cells via a Rad53-independent signaling response (Fig. 1A). To test this prediction, we compared the phosphoproteome of wild-type and rad53Δ cells using quantitative mass spectrometry and searched for Mec1-dependent signaling events triggered by checkpoint deficiency. Mec1-dependent phosphorylation was determined by crossing the dataset with previously reported phosphoproteomic analyses comparing wild-type to mec1Δ cells ^23,54^. As expected, S/T-bulky hydrophobic amino acid (ψ) motif, the Rad53 phosphorylation motif, was enriched in the set of phosphorylation events down-regulated in rad53Δ cells (Fig. S1). In contrast, phosphorylation events up-regulated in rad53Δ cells exhibited a significant enrichment of the S/T-Q motif (Fig. 1B & C), the preferential phosphorylation motif for Mec1 ^22^, indicating that loss of Rad53 triggers hyper-activation of Mec1 signaling.

**Figure 1.**
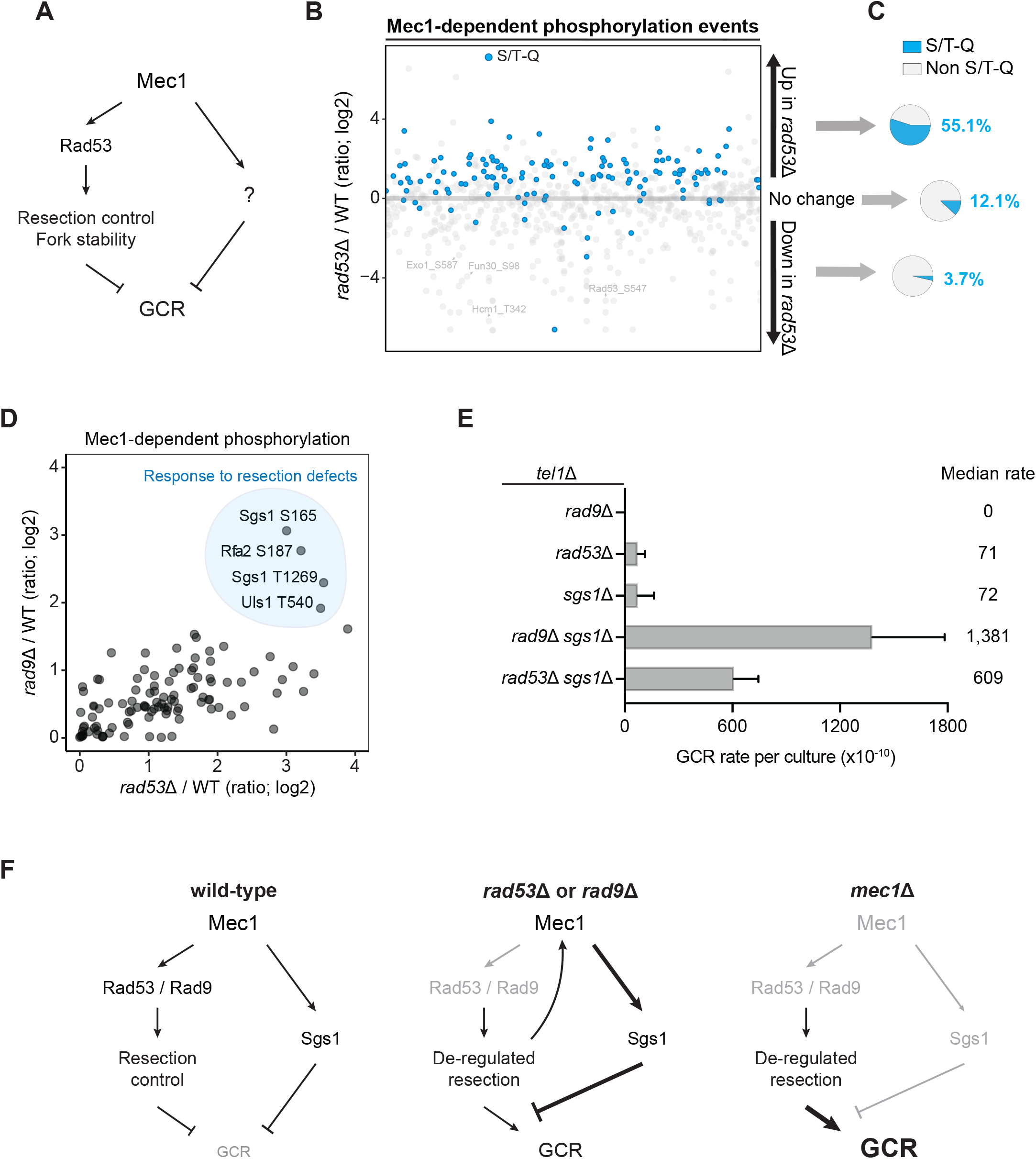
The absence of Rad53 or Rad9 induces Mec1 hyper-activation and a reliance on Sgs1 for GCR suppression. (A) Proposed model for Mec1-dependent pathways involved in GCR suppression. (B) Quantitative phosphoproteomic dataset showing the modulation of Mec1-dependent phosphorylation events in cells lacking *RAD53*, with S/T-Q consensus motifs (preferential Mec1 phosphorylation sites) indicated in blue. Cells were treated with 0.02% MMS for 2 h. (C) Pie charts showing an enrichment for S/T-Q consensus in the set of phosphorylation events upregulated in *rad53*Δ cells. (D) Quantitative phosphoproteomic data showing Mec1-dependent phosphorylation events up-regulated in both *rad9*Δ and *rad53*Δ cells. Among the most highly up-regulated sites are residues in Sgs1, Uls1, and Rfa2. (E) Measurement of GCR rates in cells with the indicated genotypes. Bars represent median values and error bars represent standard deviation from 32 individual colonies. (F) Proposed model for the involvement of Sgs1 in Mec1-dependent GCR suppression.

We previously reported that loss of Rad9, an adaptor protein that promotes Rad53 activation and the control of DNA end resection ^34–36^, triggers hyper-activation of a specialized mode of Mec1 signaling targeting proteins associated with ssDNA transactions, including Sgs1, Rfa2 and Uls1 ^54^. Interestingly, these proteins were also hyperphosphorylated in cells lacking *RAD53* (Fig. 1D), suggesting that such a response is triggered by a defect common to cells lacking *RAD9* or *RAD53*. Since Rad53 can still be active via the Mrc1 adaptor and prevent fork collapse in the absence of Rad9 ^55,56^ (Fig. S2), we reasoned that signaling hyperactivation is not triggered by replication fork collapse caused by lack of Rad53 signaling, but most likely due to increased DNA end resection, an outcome observed in both *rad53*Δ and *rad9*Δ cells ^42,57–59^. Moreover, we hypothesized that the observed hyperphosphorylation of Sgs1, a key helicase involved in multiple steps of HR and GCR suppression ^50–53,60,61^, evokes a salvage pathway that suppresses GCRs in *rad53*Δ and *rad9*Δ cells. Consistent with this model, deletion of *SGS1* displayed synergistic effects on GCR rates when combined with deletion of *RAD53* or *RAD9* (Fig. 1E). The assay was performed in a strain lacking *TEL1,* since it can partially compensate for the loss of *MEC1* in GCR ^19^. Taken together, these findings are consistent with a model whereby Mec1 suppresses GCRs through distinct pathways, one involving the control of Rad9 and Rad53, and another through the control of Sgs1 (Fig. 1F). Together with our previous report showing that DNA end hyper-resection triggers Mec1 phosphorylation of Sgs1 ^54^, our findings also suggest that upon loss of DNA end resection control via Rad53 or Rad9, the Mec1-Sgs1 pathway functions as a salvage response important to limit GCRs.

### Deregulated resection increases the demand for Mec1 control of Sgs1 in GCR suppression

To further substantiate the model proposed in Figure 1F, we investigated the importance of the phosphorylation of Sgs1 by Mec1 for GCR suppression. Sgs1 contains 9 serine or threonine residues located in the preferred motif for Mec1 phosphorylation (S/T-Q sites) (Fig. 2A). We mutated all 9 serine/threonine residues to alanine, yielding the Sgs1^9m^^ut^ mutant. Whereas expression of Sgs1^9m^^ut^ did not have a detectable effect on GCR rates in *tel1*Δ *rad9*Δ cells (Fig. 2B), we noticed that expression of Sgs1^9m^^ut^ in cells lacking *EXO1*, an exonuclease involved in DNA end resection ^10,50^, resulted in increased GCR rates. The rates of GCR accumulation caused by the expression of Sgs1^9m^^ut^ were drastically increased in cells lacking both *EXO1* and *RAD9* (Fig. 2B), further consistent with the notion that de-regulation of DNA end resection increases the demand for the Mec1-Sgs1 pathway of GCR suppression.

**Figure 2.**
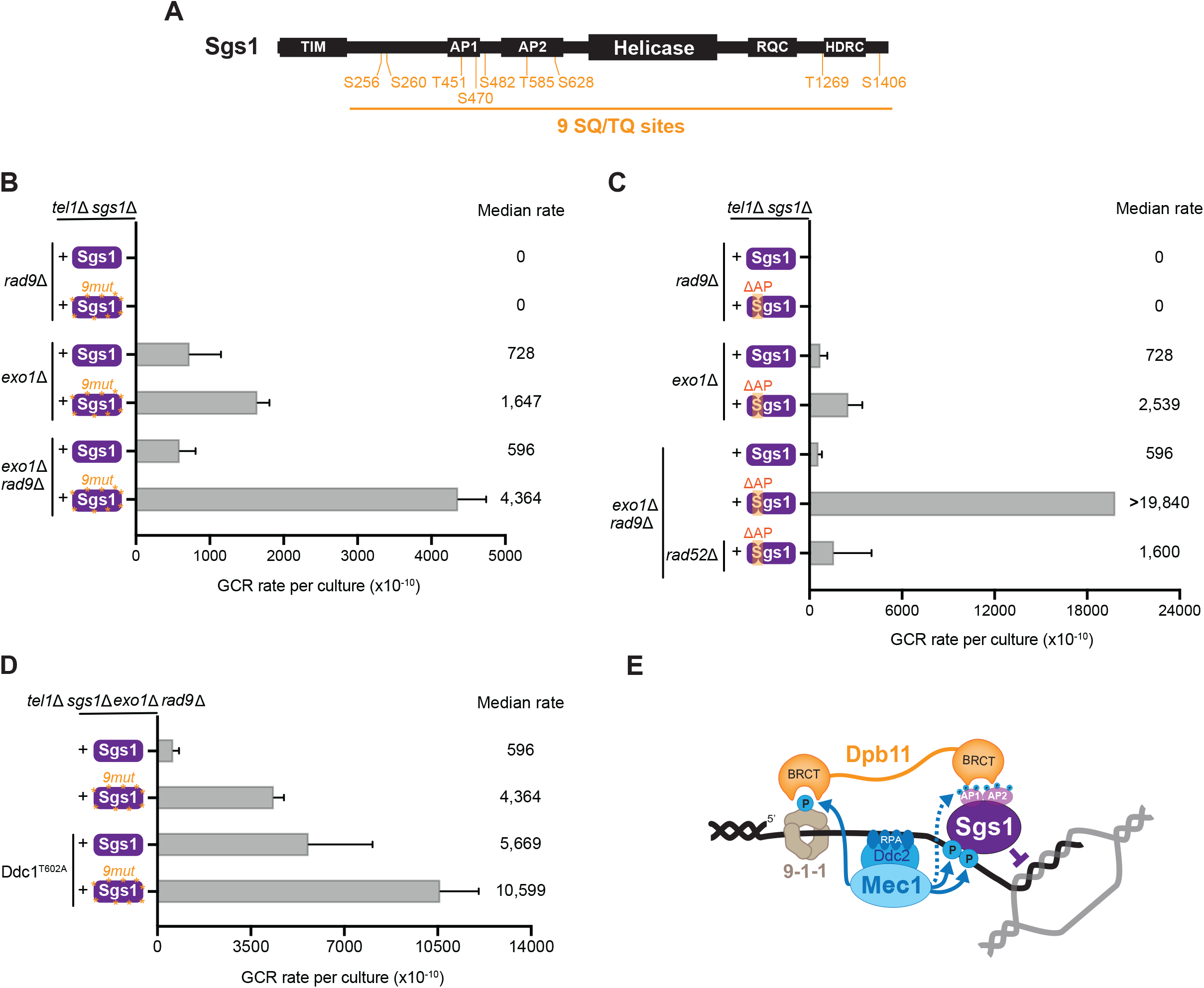
Deregulated resection increases the requirement for Mec1-dependent phosphorylation of Sgs1 in GCR suppression. (A) Schematics of Sgs1 domain architecture indicating the position of SQ/TQ sites. TIM: Top3 interacting motif; AP1/AP2: acidic patch; RQC: a region found only in the RecQ helicase family ^63^; HDRC: Helicase and RNaseD C-terminal domain. (B) Measurement of GCR rates in cells with the indicated genotypes expressing either Sgs1 or Sgs1^9m^^ut^. Bars represent median values and error bars represent standard deviation from 32 independent colonies. (C) Measurement of GCR rates in cells with the indicated genotypes expressing either Sgs1 or Sgs1^APΔ^. Bars represent median values and error bars represent standard deviation from 32 independent colonies. (D) The synergistic effect between Ddc1^T602A^ and Sgs1^9m^^ut^ on GCR suppression. Bars represent median values and error bars represent standard deviation from 32 independent colonies. (E) Speculative model for the mechanism of GCR suppression through Mec1-dependent regulation of Sgs1 that favors heteroduplex rejection. The model is based partly on our previous work showing that Mec1 mediates the recruitment of Sgs1 via the Dpb11 adaptor ^54^.

Recently we reported that Mec1 signaling promotes the interaction of Sgs1 with Dpb11 and, indirectly, to the 911 clamp, to recruit Sgs1 to DNA lesions ^54^. Consistent with our model proposing that the control of Sgs1 via Mec1 signaling is important for GCR suppression, deletion of the N-terminal acidic patches of Sgs1 (Sgs1^APΔ^ mutant) that mediate the Dpb11-Sgs1 interaction ^54^ displayed a strong increase in GCR rate in cells lacking *RAD9*, *EXO1* and *TEL1* (Fig. 2C). Sgs1^APΔ^ also failed to effectively inhibit GCRs in *rad53*Δ *exo1*Δ cells (Fig. S3B). Importantly, the high rates of GCRs observed in *tel1*Δ *rad9*Δ *exo1*Δ *sgs1*Δ cells expressing Sgs1^APΔ^ were largely dependent on Rad52 (Fig. 2C), consistent with the model that these GCRs are originating due to deregulated HR. In addition to promoting the Sgs1-Dpb11 interaction, our previous work proposed that Mec1 also promotes the recruitment of Dpb11-Sgs1 to DNA lesions by phosphorylating the Ddc1 component of the 911 clamp, which is recognized by one of the BRCT domains of Dpb11 ^54^. We therefore measured GCR rate in cells expressing the T602A mutant of Ddc1 that is not recognized by Dpb11 ^62^. As expected, expression of Ddc1^T602A^ increased GCR rates in *tel1*Δ *rad9*Δ *exo1*Δ cells (Fig. 2D), consistent with the results obtained with Sgs1^APΔ^. Surprisingly, combination of Sgs1^9m^^ut^ and Ddc1^T602A^ showed a synergistic effect on GCR suppression (Fig. 2D), suggesting that Mec1-dependent phosphorylation of Sgs1 has roles other than promoting the recruitment of Sgs1 to 911 clamp (via Dpb11). Collectively, these findings support a model in which the control of Sgs1 by Mec1 prevents GCRs driven by non-allelic HR that accumulate in cells deficient for Rad9 or Rad53 (Fig. 2E). Furthermore, the Mec1-Sgs1 pathway appears to be particularly important when the control of DNA end resection is perturbed such as in the absence of *RAD53* or *RAD9* and, especially, upon further deletion of *EXO1*. The reason for the increased importance of Sgs1 phosphorylation in the absence of *EXO1* remains unclear.

### Engineered Sgs1 recruitment suppresses GCRs in Mec1-deficient cells

Based on our proposed model (Fig. 2E), the role of Mec1 in promoting the recruitment of Sgs1 is crucial for GCR suppression, especially in cells lacking proper regulation of DNA end resection. To further test this model, we fused Sgs1 to an RPA-binding domain (RBD; amino acids 1-72 of Ddc2), with the prediction that the RBD-Sgs1 chimera would bypass the requirement of Mec1 for GCR suppression by directly recruiting Sgs1 to ssDNA at recombination intermediates, increasing heteroduplex rejection (Figs. 3A-B). Expression of RBD-Sgs1 significantly impaired break-induced replication (Fig. 3C), consistent with the expected increase in heteroduplex rejection. We have recently reported a similar effect using a fusion between Sgs1 with the BRCT domain 3/4 of Dpb11 (Dpb11^BRCT3/4^-Sgs1) ^54^, which recruits Sgs1 via recognition of the 911 clamp that is phosphorylated by Mec1, as shown in the model in Fig. 2E. We also reported that expression of Dpb11^BRCT3/4^-Sgs1 causes MMS sensitivity, presumably due to hyper-engagement of Sgs1 preventing HR-mediated DNA repair. Importantly, here we find that RBD-Sgs1 causes MMS sensitivity in both wild-type and *mec1*Δ cells, whereas Dpb11^BRCT3/4^-Sgs1 does not cause MMS sensitivity in *mec1*Δ cells (Fig. 3D). This finding is consistent with the prediction of hyper-recruitment of Sgs1 via the RBD fusion not requiring Mec1 signaling. Strikingly, RBD-Sgs1 increased Rad52 foci under both normal and MMS-treated conditions (Fig. 3E-F), which could be the consequence of increased DNA damage or slower repair process. To test whether the expression of RBD-Sgs1 generated increased DNA damage, we monitored the activation of Rad53. Expression of RBD-Sgs1 itself did not elicit Rad53 activation, nor did it impede the regular Rad53 signaling after MMS treatment (Fig. 3G). Collectively, our results suggest that RBD-Sgs1 hinders HR completion presumably by increasing heteroduplex rejection, which delays HR-mediated DNA repair, causing persistent Rad52 foci and stronger genotoxin sensitivity. Next, RBD was fused to the Sgs1^APΔ^ mutant to test the prediction that the high GCR rate observed in Sgs1^APΔ^ was caused by impaired Sgs1 recruitment and that expression of a RBD-Sgs1^APΔ^ should suppress high GCR rates. Indeed, RBD-Sgs1^APΔ^ almost eliminates GCRs in *tel1*Δ *rad9*Δ *exo1Δ sgs1Δ* cells (Fig. 3H). To further test the model that GCRs accumulate in cells lacking Mec1 due to the inability of Sgs1 to be properly recruited, we asked whether RBD-Sgs1 can suppress GCRs in Mec1-deficient cells. Since *mec1*Δ *tel1*Δ cells exhibit limited viability, we opted to use *ddc1*Δ *dna2-aa tel1*Δ cells expressing the Mec1 activation domain (MAD) of Dna2, which we have previously shown to impair Mec1 signaling and accumulate high GCR rates, while still displaying close to normal growth rates ^49^ (Fig. S4). Ectopic expression of wild-type Sgs1 or Dpb11^BRCT3/4^-Sgs1 showed similar GCR rates in *ddc1*ΔMAD *dna2-aa tel1*Δ cells, consistent with the fact that Dpb11 relies on Ddc1 for proper recruitment ^62^. In contrast, expression of RBD-Sgs1 fully suppressed GCRs, indicating that engineered Sgs1 recruitment can suppress GCRs in Mec1-deficient cells (Fig. 3I). Overexpression of Sgs1 via a strong Cyc1 promoter could also decrease the GCR rate, but the suppression was not as strong as RBD-Sgs1 (Fig. S5A-B). We further confirmed that the GCR suppression observed upon RBD-Sgs1 expression is not due to overexpression of the fusion protein since the RBD-Sgs1 fusion was in fact less abundant than Sgs1 (Fig. S5C&D). Expression of an Ddc1^T602A^-Sgs1 fusion (causing constitutive recruitment to the 911 clamp) could also efficiently suppress GCRs in *dna2-aa ddc1Δ tel1Δ* cells (Fig. S6), further supporting the model that in cells lacking Mec1, GCRs accumulate due to the inability of Sgs1 to be properly recruited.

**Figure 3.**
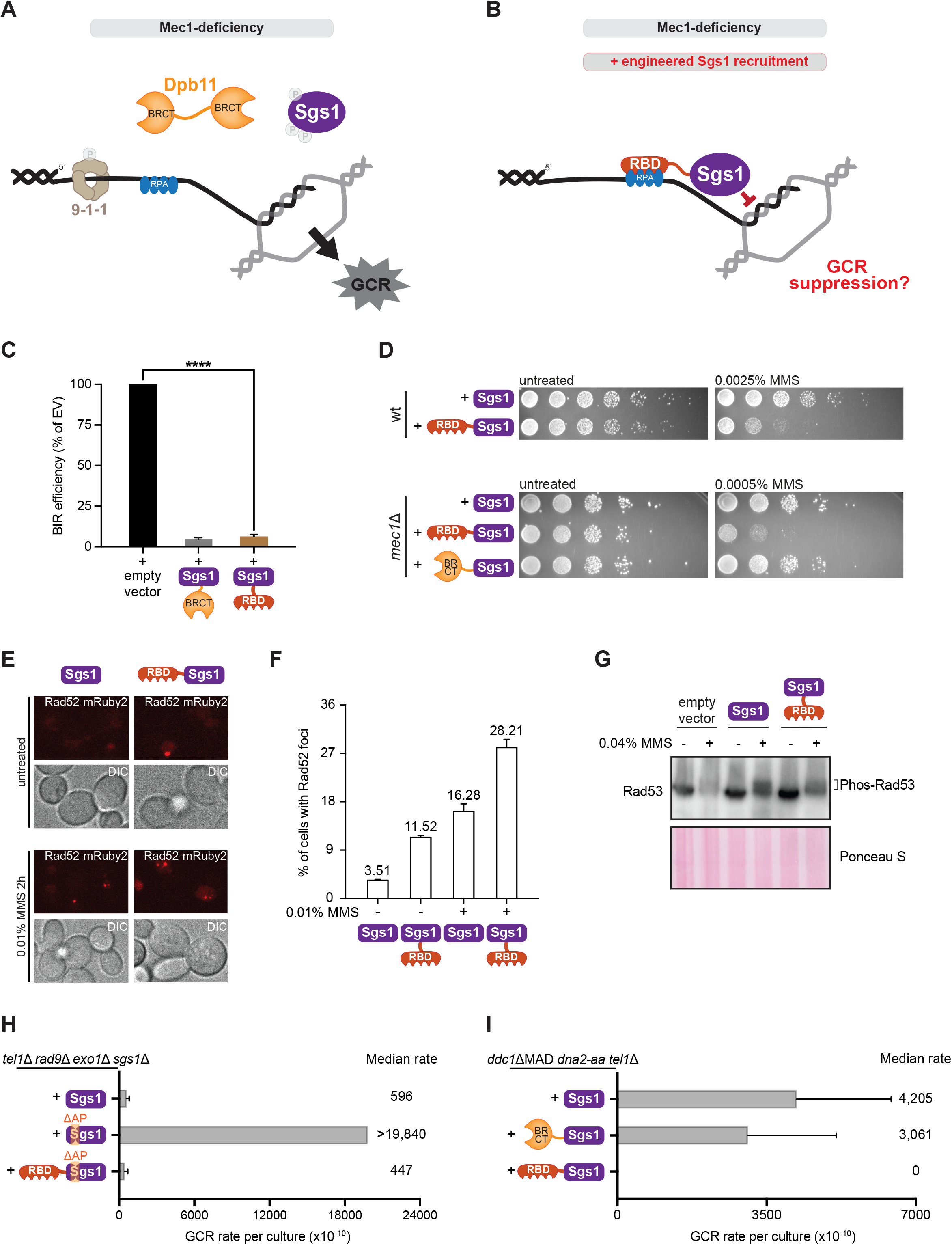
Engineered Sgs1 recruitment suppresses GCRs in Mec1-deficient cells. (A) Schematics illustrating how the lack of Mec1-mediated Sgs1 recruitment leads to increased GCRs. (B) Schematics depicting the rationale for designing an RBD-Sgs1 chimera for recruitment of Sgs1 independently of Mec1 signaling. (C) Measurement of BIR efficiency in cells carrying an empty vector or expressing different Sgs1 chimeras. Bars represent mean values and error bars represent standard deviation from three replicate experiments. P value was calculated with a two-tailed, unpaired t-test. ****P ≤ 0.0001. (D) Dilution assay for monitoring MMS sensitivity of wild-type or *mec1*Δ cells expressing RBD-Sgs1 or Dpb11^BRCT3/4^-Sgs1. (E) Representative image of Rad52 foci in cells expressing Sgs1 or RBD-Sgs1 untreated or treated with 0.01% MMS for 2 h. (F) Quantification of percentages of cells with Rad52 foci from E. Over 150 cells were scored per replicate. Bars represent mean values and error bars represent standard error of the mean from three replicate experiments. (G) Western blot showing Rad53 mobility shift induced by MMS in cells expressing either Sgs1 or RBD-Sgs1. (H) Measurement of GCR rates in *tel1*Δ *rad9*Δ *exo1*Δ *sgs1*Δ cells expressing either Sgs1, Sgs1^APΔ^, or RBD-Sgs1^APΔ^. Bars represent median values and error bars represent standard deviation from 32 independent colonies. (I) Measurement of GCR rates in *ddc1*ΔMAD *dna2-aa tel1*Δ cells expressing Sgs1, Dpb11^BRCT3/4^-Sgs1, or RBD-Sgs1. Bars represent median values and error bars represent standard deviation from 32 independent colonies.

Sgs1 is a large multi-domain protein (Fig. S7A). To define the critical regions required for the GCR suppressive function of Sgs1 when fused to RBD, we generated several mutations and truncations in Sgs1 and monitored GCR rates. Removal of the Top3 interacting motif (TIM, 1-158aa) did not affect the function of the chimera (Fig. 4A), indicating that the ability of Sgs1 to bind to Top3 was not necessary for GCR suppression when Sgs1 was hyper-recruited. Chimeras with either helicase-defective mutation (hd, Sgs1^K706A^) or deletion of the RQC domain (1081-1195aa, a region found only in the RecQ helicase family ^63^) lost the ability to suppress GCRs (Fig. 4A), showing that the helicase activity of Sgs1 is essential for GCR suppression. Loss of the Helicase and RNaseD C-terminal domain (HRDC, 1271-1351aa), which is involved in DNA binding ^64^, caused no change in GCR rate (Fig. 4A). Similar results were obtained using *ddc1*Δ *tel1*Δ *rad53*Δ cells, among which Mec1 signaling is partially disrupted (Fig. S8).

**Figure 4.**
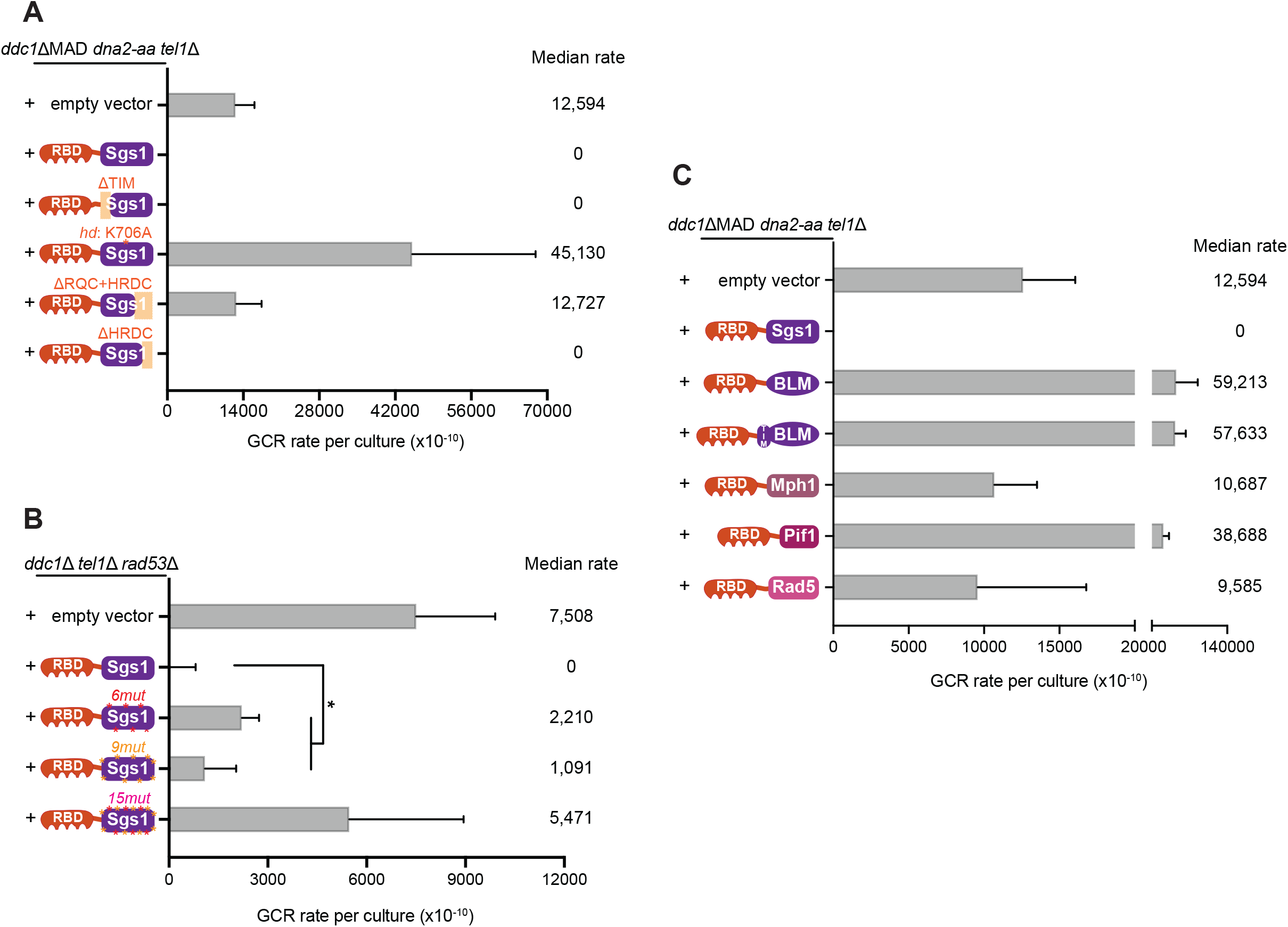
GCR suppression through the RBD-Sgs1 chimera requires Sgs1 helicase activity and Sgs1 phosphorylation. (A) Measurement of GCR rates in *ddc1*ΔMAD *dna2-aa tel1*Δ cells expressing RBD fused to wild-type Sgs1 or truncations of Sgs1 (*hd*: helicase-dead; see legend in 2A for description of domains). Bars represent median values and error bars represent standard deviation from 32 independent colonies. (B) Measurement of GCR rates in *ddc1*ΔMAD *dna2-aa tel1*Δ cells expressing RBD fused to wild-type Sgs1 or Sgs1 containing phospho-site mutations (*6mut*: mutation of 6 sites including 4 SP/TP sites; *9mut*: mutation of 9 SQ/TQ sites; *15mut*: combination of *6mut* and *9mut* mutations). Bars represent median values and error bars represent standard deviation from 32 independent colonies. (C) Measurement of GCR rates in *ddc1*ΔMAD *dna2-aa tel1*Δ cells expressing RBD fused to yeast DNA helicases Sgs1, Mph1, Pif1 or Rad5, or fused to BLM, the human ortholog of Sgs1. Bars represent median values and error bars represent standard deviation from 16 independent colonies.

Since Mec1 phosphorylation of Sgs1 has recruitment-independent roles (Fig. 2D), we predicted that RBD-Sgs1^9m^^ut^ would partially weaken GCR suppression. We used *ddc1*Δ *tel1*Δ *rad53*Δ cells to test this hypothesis because in this strain Sgs1 recruitment via 911-Dpb11 is impaired while Mec1 signaling is still functional via Dna2-mediated activation. As expected, loss of Mec1 phosphorylation impaired the suppression of GCRs; however, the change was modest (Fig. 4B). When we introduced serine-to-alanine mutations at other 6 positions, containing 4 putative CDK phosphorylation sites (Fig. S7B), serine-proline motifs, we also observed a modest increase in GCR rate (Fig. 4B). Strikingly, impairing both Mec1 and CDK phosphorylation motifs in Sgs1 by combining all 15 mutations (RBD-Sgs1^15m^^ut^) led to synergistic effects (Fig. 4B), suggesting that both Mec1 and CDK promote Sgs1’s function in GCR suppression.

Next, we asked whether GCRs can be suppressed by other helicases when fused to RBD, or if Sgs1 has unique properties that confer its GCR suppressive function. We fused RBD to other yeast helicases involved in recombination, including Mph1, Pif1 and Rad5 ^61^, and found that none of them had the ability to prevent GCR accumulation (Fig. 4C). Fusing RBD to BLM, the human ortholog of Sgs1, also could not inhibit GCRs and even showed a higher GCR rate, similar to helicase-dead Sgs1 (Fig. 4A). Addition of the Top3 interacting motif of Sgs1 to RBD-BLM did not alter GCR rates (Fig. 4C). Taken together, our results showed that engineered Sgs1 recruitment can effectively suppress GCRs, and this function is highly specific to Sgs1.

### Engineered Sgs1 recruitment suppresses HR-driven GCRs and eliminates D-loop formation

Since Sgs1 functions at multiple steps in HR, including DNA end resection, heteroduplex rejection and double Holliday junction (dHj) dissolution ^50–53^, we sought to determine which step in HR is impacted by RBD-Sgs1 and likely contributing to the suppression of GCRs. Defects in heteroduplex rejection can give rise to non-allelic HR events ^65^, while inefficient dHj dissolution can increase the occurrence of crossovers ^51^. Defects in both of these processes can induce chromosomal rearrangements. Multiple lines of evidence support the hypothesis that RBD fusion enhances the capacity of Sgs1 to reject heteroduplexes, thereby preventing GCRs driven by non-allelic HR. First, we showed that removal of the TIM of Sgs1 does not affect the ability of RBD-Sgs1 to suppress GCR (Fig. 4A), excluding the requirement Top3’s role in strand passage for joint molecule dissolution ^53,66^. Second, RBD-Sgs1 could still suppress GCRs in *rad51*Δ cells (Fig. 5A), where Holliday junctions do not form although Rad52-dependent single-strand annealing can still be used ^67^, which would necessitate heteroduplex rejection for HR quality control. In this context, if RBD-Sgs1 suppresses GCRs by promoting heteroduplex rejection, RBD-Sgs1 should fail to suppress GCRs in *rad52*Δ cells. Indeed, in the absence of Rad52, the effect of RBD-Sgs1 expression was comparable to the expression of Sgs1 (Fig. 5B). To directly monitor the effect of RBD-Sgs1 in heteroduplex rejection, we performed the displacement loop (D-loop) capture (DLC) assay (Fig. 5C and ^61^), where stronger rejection of heteroduplexes results in decreased DLC signal. Consistent with the hypothesis that engineered Sgs1 recruitment suppresses GCR by promoting heteroduplex rejection, both Dpb11^BRCT3/4^-Sgs1 and RBD-Sgs1 expression nearly eliminates the D-loop formation (Fig. 5D & Fig. S9A). Taken together, our results showed that engineered Sgs1 recruitment suppresses HR-driven GCRs through enhanced heteroduplex rejection.

**Figure 5.**
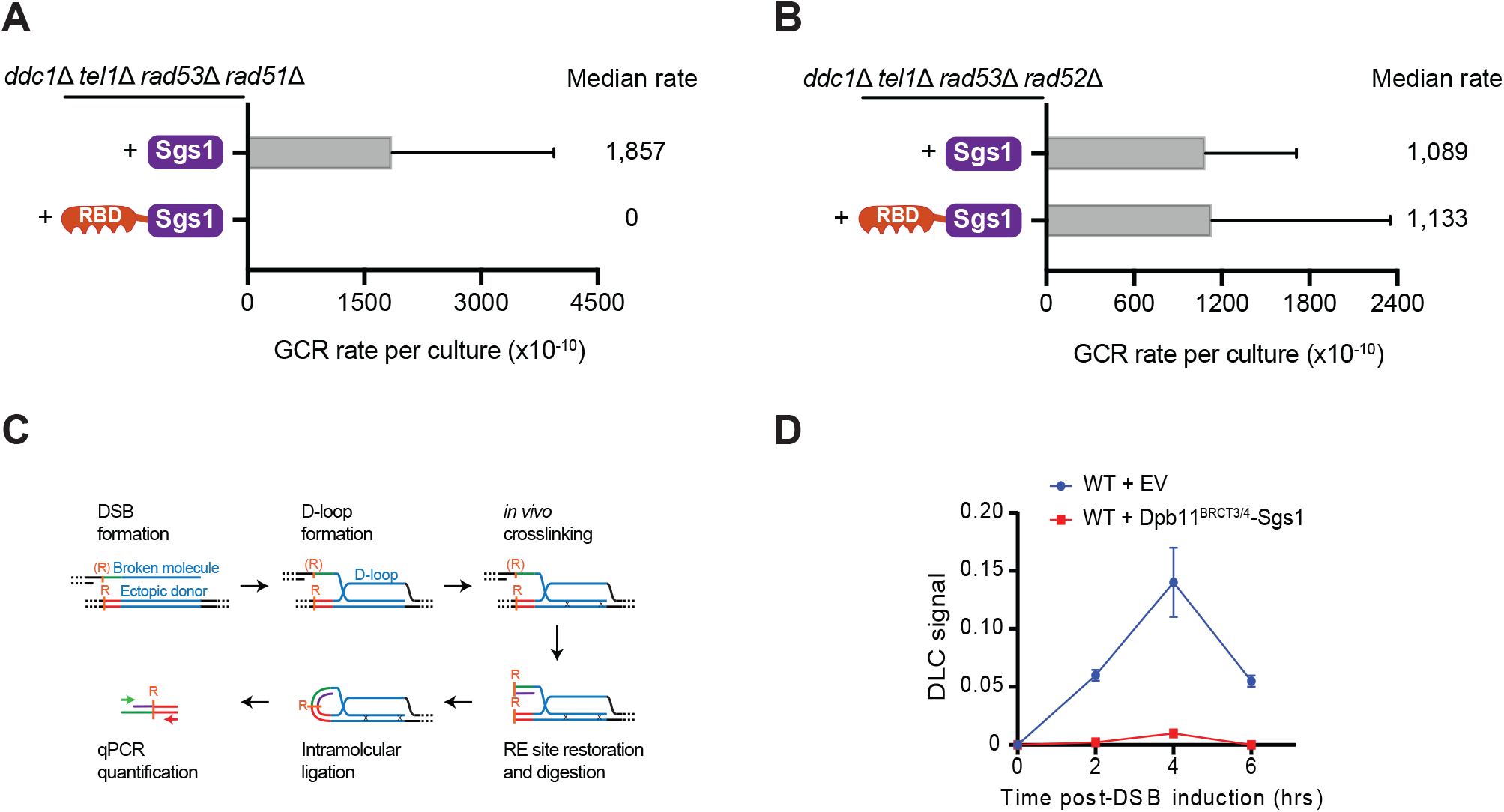
Engineered Sgs1 recruitment via RBD-Sgs1 chimera suppresses HR-driven GCRs and eliminates D-loop formation. (A) Measurement of GCR rates in *ddc1*Δ *tel1*Δ *rad53*Δ *rad51*Δ cells expressing either Sgs1 or RBD-Sgs1. Bars represent median values and error bars represent standard deviation from 32 independent colonies. (B) Measurement of GCR rates in *ddc1*Δ *tel1*Δ *rad53*Δ *rad51*Δ cells expressing either Sgs1 or RBD-Sgs1. Bars represent median values and error bars represent standard deviation from 32 independent colonies. (C) Schematic representation of the D-loop capture (DLC) assay ^61^. (D) DLC signal in cells carrying an empty vector or expressing the DPB11^BRCT3/4^-Sgs1 chimera. Error bars represent SEM of two replicate experiments.

## Discussion

Over 20 years ago, foundational work by the group of Richard Kolodner revealed elevated rates of GCRs in *mec1*Δ *tel1*Δ cells ^19^. Surprisingly, their work also showed that the ability of Mec1 and Tel1 to suppress GCRs is largely independent of their canonical role in activating the DNA damage checkpoint ^19,49^. The detailed mechanism by which Mec1 and Tel1 suppress GCRs has remained elusive, representing a major gap in our understanding of kinase-mediated genome maintenance mechanisms. Here we focused on the GCR suppressive function of Mec1 and found that *mec1*Δ cells accumulate GCRs that are driven by deregulated HR. Moreover, we revealed that higher GCR rates are caused by compounding effects from the combined loss of DNA damage checkpoint and the control of Sgs1 (Fig. 6). Our findings show that, upon loss of DNA damage checkpoint signaling and resection control, a Mec1-Sgs1 salvage pathway limits GCR accumulation. We propose that this salvage pathway increases heteroduplex rejection, functioning as a boosted HR quality control mechanism that limits non-allelic recombination.

**Figure 6.**
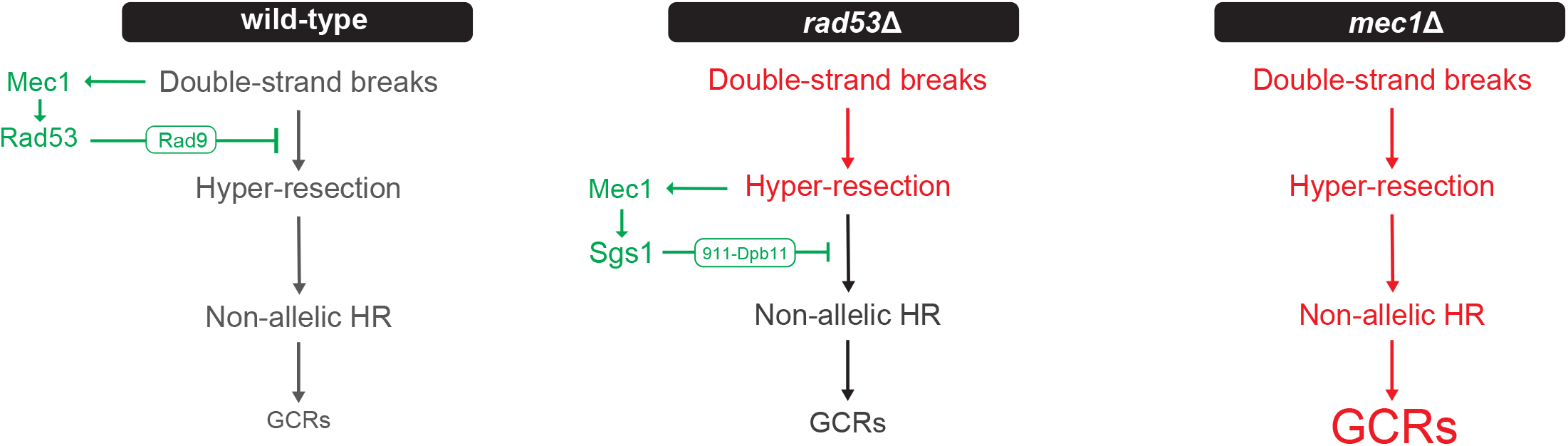
Model for GCR suppression via multi-step control of HR by Mec1. Upon DSB and initial end resection, Mec1 is recruited to RPA-ssDNA to promote the Rad9-Rad53 signaling axis that restrains long range resection. This anti-resection function of Mec1 protects DNA ends from extensive nucleolytic processing, thereby reducing the chance of non-allelic HR and preventing GCRs. In cells lacking *RAD9* or *RAD53*, DNA ends undergo hyper-resection, which activates a mode of Mec1 signaling leading to Sgs1 phosphorylation, and its recruitment to lesion sites via the 911-Dpb11 complex. This recruitment results in the inhibition of non-allelic HR through heteroduplex rejection, thereby suppressing GCRs. Mec1 phosphorylation of Sgs1 also suppresses GCRs through, yet unknown, recruitment-independent mechanisms. In contrast, *mec1*Δ cells fail to restrain resection and also lack the Mec1-Sgs1 salvage pathway (impaired HR quality control), leading to a dramatic increase of non-allelic HR driven GCRs.

Quantitative phosphoproteomic analysis of *rad53*Δ or *rad9*Δ cells showed that these mutants display increased Mec1 signaling directed towards a selective group of proteins involved in ssDNA transactions. In particular, the hyperphosphorylation of the Sgs1 helicase in these strains promotes its recruitment to DNA lesion sites via the association with the 911-Dpb11 complex. The discovery of novel modes of Mec1/ATR signaling upon loss of checkpoint reveals the multi-faceted action and complex regulation of this kinase. Since *rad9*Δ cells do not suffer the drastic replication fork collapse phenotype observed in *rad53*Δ cells (Fig. S2), we favor the model that the hyper-activation of Mec1 observed in both *rad53*Δ or *rad9*Δ cells is caused by deregulated resection. Notably, the lack of Rad53 has been shown to impair Rad9’s role in counteracting resection ^36,59^, consistent with both *rad53*Δ or *rad9*Δ cells sharing a similar defect in resection control. Exactly how deregulated resection promotes Mec1 signaling is still unclear. One possibility is that faster rates of resection, or imbalanced engagement of resection nucleases Exo1 and Dna2, causes abnormal exposure of ssDNA that is sensed by Mec1. Since increased exposure of ssDNA is expected to increase non-allelic recombination, an interesting implication of our model is that the signal for Mec1 activation is the actual driver of GCR events, implying that Mec1 signaling serves as a rheostat to increase heteroduplex rejection and HR quality control. We propose that tightly controlling heteroduplex rejection in a context-dependent manner, and not overstimulating it when not needed, is crucial to make sure HR can be properly utilized for DNA repair transactions, such as template switching, when needed. Moreover, our findings, and previous reports ^68^, highlight an important role for Exo1 in preventing GCRs and that *exo1*Δ cells have an increased demand for Sgs1 regulation. Whereas Exo1 and Dna2-Sgs1 are involved in extensive resection ^10,50^, Rad9 was reported to prevent hyper-resection by Sgs1 ^69^, with faster resection in *rad9*Δ cells being mainly dependent on Sgs1 ^70^. Thus, Exo1 may play an important role in competing with Dna2-Sgs1, which may ensure proper resection.

In the context of the RBD-Sgs1 chimera, it is surprising that the Sgs1-Top3 interaction is not required for GCR suppression. Previous studies have shown that it is the Top3 activity that reverses D-loop and that the helicase activity of Sgs1 is not required for D-loop disruption ^61,71^. However, our findings show that Sgs1’s helicase activity is essential for GCR suppression while the Sgs1-Top3 interaction is not required. A key difference is that previous studies used assays with homologous D-loops, while GCRs in our systems are expected to mainly arise from homeologous recombination. Therefore, our results suggest that the rejection of homeologous heteroduplexes requires the helicase activity of Sgs1. It is tempting to speculate that, in this context, the helicase activity of Sgs1 is linked to the mismatch recognition complex Msh2-Msh6 to recognize homeologous heteroduplexes. In support of this idea, Sgs1 and Msh6 were shown to play similarly important roles in heteroduplex rejection ^52,72,73^. Additional lines of circumstantial evidence from our work are consistent with the model that Sgs1 requires its helicase activity to favor the rejection of heteroduplexes with homeologous sequences. For example, we found that the expression of the RBD-Sgs1^hd^ chimera induces a dramatic increase in GCR rates, which could be caused by its ability to effectively disrupt homologous D-loops while failing to disrupt homeologous D-loops, therefore mainly driving repair based on non-allelic homeologous sequences. Moreover, our results also show that fusing other yeast helicases to the RBD domain does not result in appreciable GCR suppression as seen with the RBD-Sgs1 chimera, potentially due to the fact that these other helicases do not interact with Msh6 and/or do not have their helicase activity coupled to mismatch recognition. Notably, expression of RBD-BLM in Mec1-deficient cells generated more GCRs, with values similar to that of the RBD-Sgs1^hd^. Though BLM was shown to interact with human MSH6 both in vivo and in vitro ^74^, it may not be able to interact with yeast Msh6. Thus, similar to RBD-Sgs1^hd^, RBD-BLM may be able to disrupt homologous D-loops but fails to efficiently disrupt D-loops between homeologous sequences, resulting in more non-allelic HR events.

In the future, it will be important to investigate how Mec1 phosphorylation modulates the helicase activity of Sgs1, and how the phosphorylation events alter the ability of Sgs1 to reject heteroduplexes with homeologous sequences. Our results show that RBD-Sgs1 requires phosphorylation at both CDK and Mec1 sites to efficiently prevent GCR accumulation. Notably, CDK phosphorylation of Sgs1 has been shown to stimulate DNA unwinding ^75^. Our finding that cells expressing Sgs1^6m^^ut^ (CDK sites mutated) display increased MMS sensitivity whereas cells expressing Sgs1^9m^^ut^ (S/T-Q sites mutated) do not exhibit genotoxin sensitivity (Fig. S10) suggests that Mec1 signaling plays a more specialized role in the regulation of Sgs1 action and HR quality control, perhaps by fine-tuning the stringency of the detection of homeologous heteroduplexes.

Although this work addresses how Mec1 prevents non-allelic HR driven GCRs, it is worth mentioning that GCRs can also arise by HR-independent pathways, such as *de novo* telomere addition and non-homologous end joining (NHEJ) ^13,17^. While in *rad51*Δ cells, heteroduplex rejection can still occur during Rad52-mediated SSA ^52^ with RBD-Sgs1 still disrupting D-loops and inhibiting non-allelic HR, in *rad52*Δ cells GCRs are expected to be caused by non-HR pathways such as NHEJ, with RBD-Sgs1 failing to suppress GCRs.

Mec1 is expected to suppress GCRs through additional mechanisms that do not require Sgs1, as evidenced by a comparison of GCR rates in different mutant strains. For example, the GCR rates of *tel1*Δ *rad9*Δ *exo1*Δ Sgs1^9m^^ut^ Ddc1^T602A^ (∼11,000) are significantly lower compared to that of *mec1*Δ *tel1*Δ (∼45,000). Consistent with Sgs1-independent roles for Mec1 in GCR suppression, our phosphoproteomic analysis revealed that loss of *RAD9* or *RAD53* induces phosphorylation of other proteins with roles in ssDNA-associated transactions, such as Rfa2 and the ubiquitin ligase and DNA translocase Uls1. Further dissecting the roles of these, and potentially other, Mec1 phosphorylation events induced in *rad9*Δ cells should shed light into additional GCR suppressing mechanisms controlled by Mec1. Moreover, it will be important to define the role of Tel1 in limiting GCR accumulation upon loss of Mec1. One possibility is that DSBs accumulate in *mec1*Δ cells due to increased fork collapse, and that Tel1 is required to properly repair these breaks and prevent them from engaging in deleterious DNA transactions that cause GCRs.

Whereas yeast offers a robust and much simplified system to dissect mechanisms of GCR suppression, we envision that our findings may contribute to better understanding GCR suppression mechanisms in mammals. For example, exploring how mammalian cells respond to de-regulated resection may uncover similar salvage pathways involved in heteroduplex rejection control as the Mec1-Sgs1 pathway identified here. Interestingly, BLM has been shown to interact with TOPBP1, the ortholog of Dpb11, although the interaction is not dependent on ATR ^76–78^. Nevertheless, BLM is phosphorylated by ATR ^79^, which could have an effect on BLM’s function in heteroduplex rejection. It is also possible that ATR may respond to de-regulated resection in a more complex manner than Mec1 does in yeast, involving a larger set of substrates and GCR suppression mechanisms. Moreover, it is also possible that ATR-independent responses are triggered upon de-regulated resection and actively control heteroduplex rejection to limit genetic instability. In summary, exploring the response to de-regulated resection in mammals may open new directions to understand mechanisms of genome maintenance.

## Materials and Methods

### Yeast strains

A complete list of yeast strains used in this study can be found in Supplemental Table S2. The strain background for all yeast used in this study is S288C, unless indicated. Gene deletions were performed using standard PCR-based strategy to amplify resistance cassettes with flanking sequences homologous to the target gene. All endogenous deletions were verified by PCR. Plasmids in this study are listed in Supplemental Table S3 and are available upon request. Yeast strains were grown at 30 °C in a shaker at 220 rpm. For strains with endogenous deletion, YEPD media were used. For strains carrying plasmids, the corresponding synthetic dropout media were used. For SILAC experiments, yeast strains were grown in -Arg -Lys media supplemented with either isotopically normal arginine and lysine (“light” media) or the ^13^C^15^N isotopologue (“heavy” media). Excess proline was added to SILAC media at a concentration of 80 mg/L to prevent conversion of arginine to proline.

### Western blots

50 ml of yeast were grown in appropriate media to mid-log phase and treated as described in the figure legend. Cells were pelleted at 1,000 rcf and washed with TE buffer (pH 8.0) containing 1 mM PMSF. Pellets were lysed by bead beating with 0.5-mm glass beads for three cycles of 10 min with 1 min rest time between cycles at 4°C in lysis buffer (150 mM NaCl, 50 mM Tris pH 8.0, 5 mM EDTA, 0.2% Tergitol type NP-40) supplemented with complete EDTA-free protease inhibitor cocktail (Roche), 1 mM PMSF, 5 mM sodium fluoride, and 10 mM b-glycerophosphate. Concentration normalization was performed via the Bradford assay. Lysates were boiled in Laemmli buffer and electrophoresed on a 10% SDS–PAGE gel. Proteins were then transferred wet onto a PVDF membrane and incubated with antibody. Signal detection was performed using HRP-coupled secondary antibodies, imaged with BioRad ChemiDoc.

### Phosphoproteomics

For phosphoproteomic experiments, 150 ml of yeast were grown in “heavy” or “light” SILAC media to mid-log phase and treated with 0.04% MMS for 2h. Cells were pelleted and lysed as described for western blots above. Protein digestion, phosphoenrichment and following MS data analysis were performed as described in 50.

### Immunoprecipitation–mass spectrometry (IP-MS)

For IP-MS experiment, 150 ml of yeast were grown in “heavy” or “light” SILAC media to mid-log phase. Cells were pelleted and lysed as described for western blots above. Around 5 mg of lysate per sample was incubated with antibody-conjugated agarose resin (Anti-c-Myc, Sigma) for 3h at 4°C. Resin was washed 4 times in the lysis buffer. Proteins were eluted by heating at 65°C with elution buffer (1% SDS, 100 mM Tris pH 8.0) for 15 min. MS samples preparation were performed as described in 50.

### GCR assays

All GCR assays were performed with yeast freshly streaked from frozen glycerol stocks or new transformations. Plates were incubated at 30°C for 3-4 days to get visible colonies. Individual colonies with similar sizes were picked and transferred to 2 ml of culture (YPD for strains with integrated genetic modification, -Leu media for strains with pRS415 plasmids). After 48h, ∼10 million cells were spun down, washed with 400μl of autoclaved ddH_2_O, resuspended in 150∼200μl of autoclaved ddH_2_O and spotted onto plates containing 5-FOA and canavanine ^80^. Fewer cells were used when strains have extremely high GCR rates, e.g., *exo1*Δ *sgs1*Δ. In parallel with each GCR experiment, multiple cultures (usually 4 in this study) were randomly chosen and serially diluted (for YPD, 2×10^6; for -Leu, 5×10^4) and plated onto YPD plates to determine the average population viability. After 4 days, the number of 5-FOA- and canavanine-resistant colonies in a spot was counted. The number of GCR events in a culture was calculated using the equation m[1.24 + ln(m)] − r = 0, where r is the number of 5-FOA- and canavanine-resistant colonies in a spot, and m is the estimated number of GCR events^80^. GCR rate was then calculated by dividing the number of GCR events per culture by the average population viability. For each GCR experiment, at least 16 independent colonies were picked and 2 independent strains with the same genotype were used.

### D-loop capture assay

For D-loop capture experiments, all strains were in the W303 RAD5 background. They contain a copy of the GAL1/10 driven HO endonuclease gene at the TRP1 locus on chr. IV. A point mutation inactivates the HO cut-site at the mating-type locus (MAT) on chr. III (*MATa-inc*). The DSB-inducible construct contains the 117 bp HO cut-site, a 2,086 bp-long homology A sequence (+4 to +2090 of the LYS2 gene), and a 327 bp fragment of the PhiX174 genome flanked by multiple restriction sites ^61^. D-loop capture assay was performed as previously reported ^61,81^, with the following modifications: zymolyase lysed cells were proceeded immediately to the restriction digestion, ligation and DNA purification step after hybridization with oligonucleotides as described previously ^82^.

### Microscopy analysis

For Rad52 foci analysis, cells were grown at 30°C in synthetic complete media (for *rad9*Δ and *rad53*Δ microscopy) or -Leu media (for Sgs1 rejector microscopy) until OD_600_ reaches 0.2, and 0.01% MMS was added to the culture for 2 h if mentioned. Next, 200μl of culture was transferred to 4-chamber glass bottom dishes (Cellvis), which were pre-treated with 0.5 mg/ml concanavalin A (Sigma). After 5 min of fixation, liquid was aspirated, and cells were washed with 200μl of autoclaved ddH_2_O. 1 ml of requisite media was added to keep cells alive during imaging. Over 150 cells were scored for each replicate. Images were acquired at room temperature using a spinning-disc confocal microscope (CSU-X; Yokogawa Electric Corporation and Intelligent Imaging Innovations) on an inverted microscope (DMI600B; Leica Biosystems) with a 100×, 1.46 NA objective lens and an electron-multiplying charge-coupled device camera (QuantEM; Photometrics). 488nm laser lines were used for the detection of mRuby-tagged Rad52 in yeast cells. SlideBook software (Intelligent Imaging Innovations) was used to obtain Z stack images. Maximum intensity projections were created in the Slidebook software for foci number analysis.

### Dilution assays

For dilution assays, 3 ml of yeast culture was grown to saturation at 30°C. Then, 1 OD_600_ equivalent of the saturated culture was serially diluted (10-fold serial dilutions were used unless noted) in a 96-well plate with autoclaved ddH_2_O and spotted onto agar plates using a bolt pinner. Plates were incubated at 30°C for 2 days before imaging.

## Acknowledgements

We thank Beatriz S. Almeida for technical support; we thank members of the Smolka Lab for valuable discussions related to this work. This work is supported by grants from the National Institute of Health, R35GM141159 to M.B.S., and R01GM58015 and R01GM137751 to W.D.H. S.H.H. was partially supported by a fellowship from the Academia Sinica, Taiwan.

## Author contribution

B.X and M.B.S conceptualized the project, designed the experiments, interpreted the data, and wrote the paper. B.X, E.J.S, and M.M.W performed experiments. B.X, E.J.S, and M.M.W analyzed the phosphoproteomic data. S.H.H performed the D-loop capture experiments and W.D.H. helped design the experiment and interpret the data. All authors contributed to revising the manuscript.

---

**Figure S1, related to figure 1.**
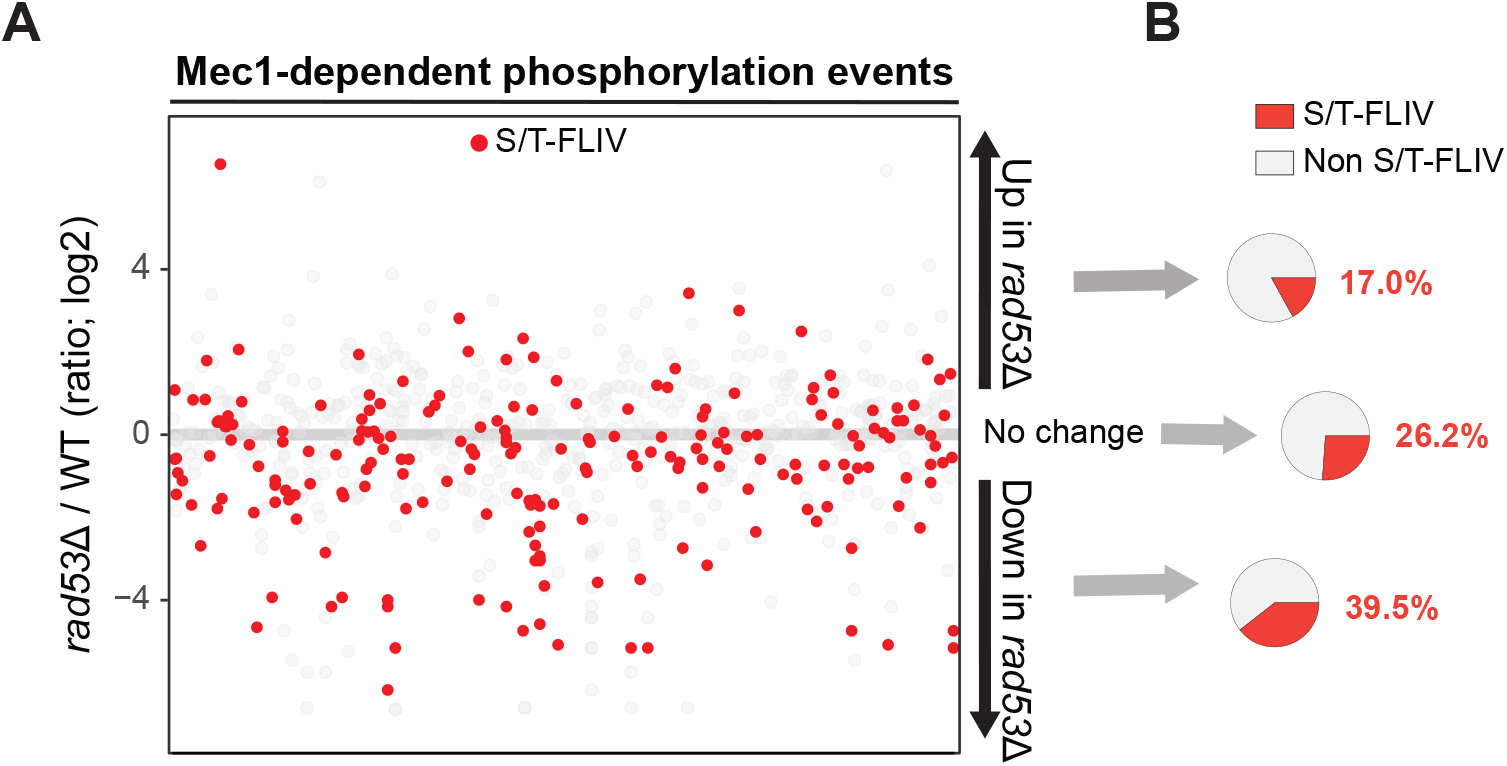
Rad53-dependent signaling enriched for S/T-FLIV motifs is down-regulated in *rad53*Δ cells. (A) Quantitative phosphoproteomic data showing the modulation of Mec1-dependent phosphorylation events in cells lacking RAD53, with S/T-FLIV consensus motifs (preferential Rad53 phosphorylation sites) indicated in red. Cells were treated with MMS for 2 h. (B) Pie chart showing that S/T-FLIV consensus phosphorylation events are downregulated in *rad53*Δ cells.

**Figure S2, related to figure 1.**
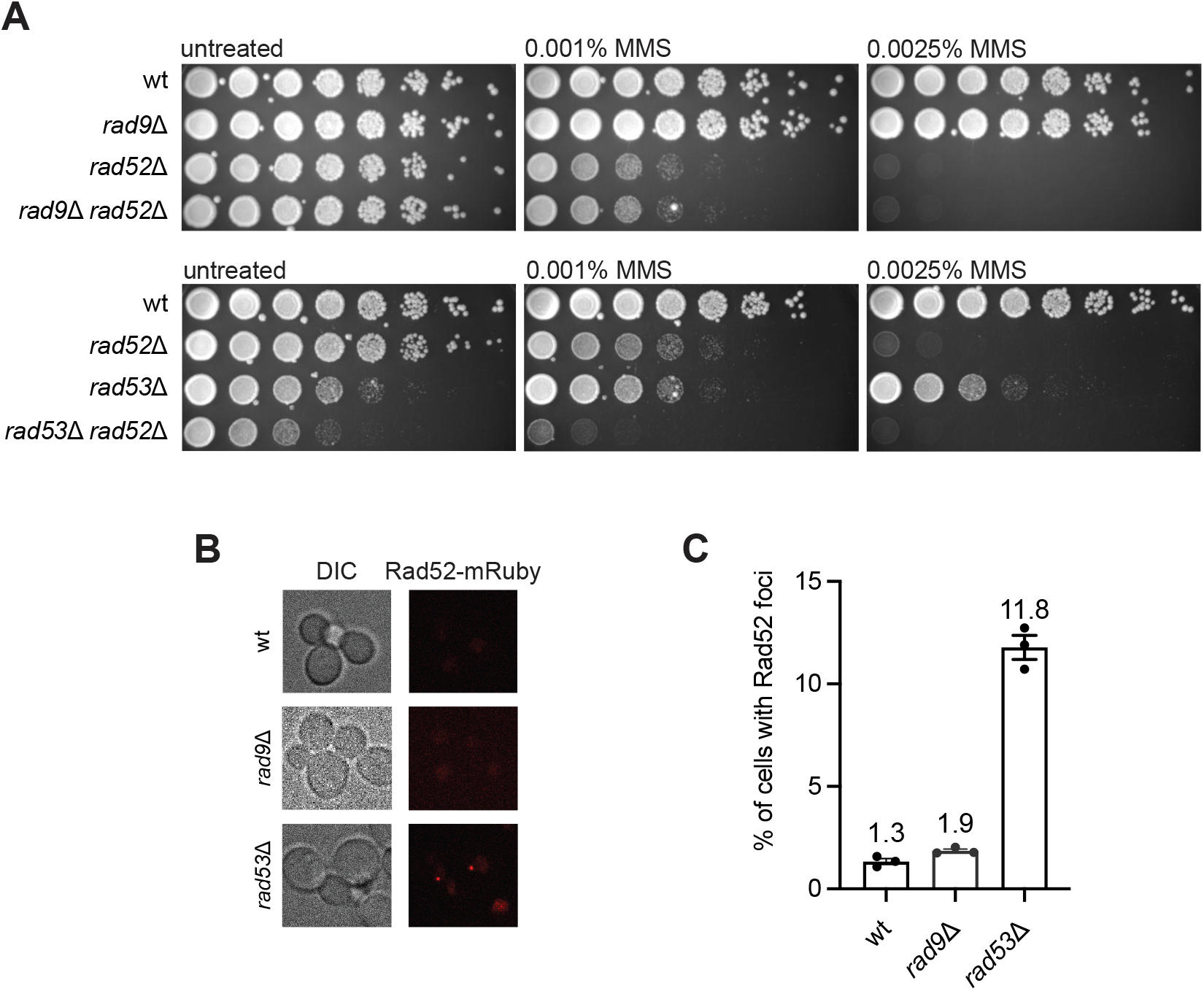
*rad53*Δ cells, but not *rad9*Δ cells, display increased demand for HR. (A) Dilution assay of cells with indicated genotype in the presence of MMS. (B) Representative image of Rad52 foci in cells with indicated genotype under untreated condition. (C) Quantification of percentages of cells with Rad52 foci. Over 150 cells were scored per replicate. Bars represent mean values and error bars represent standard error of the mean from three replicate experiments.

**Figure S3, related to figure 2.**
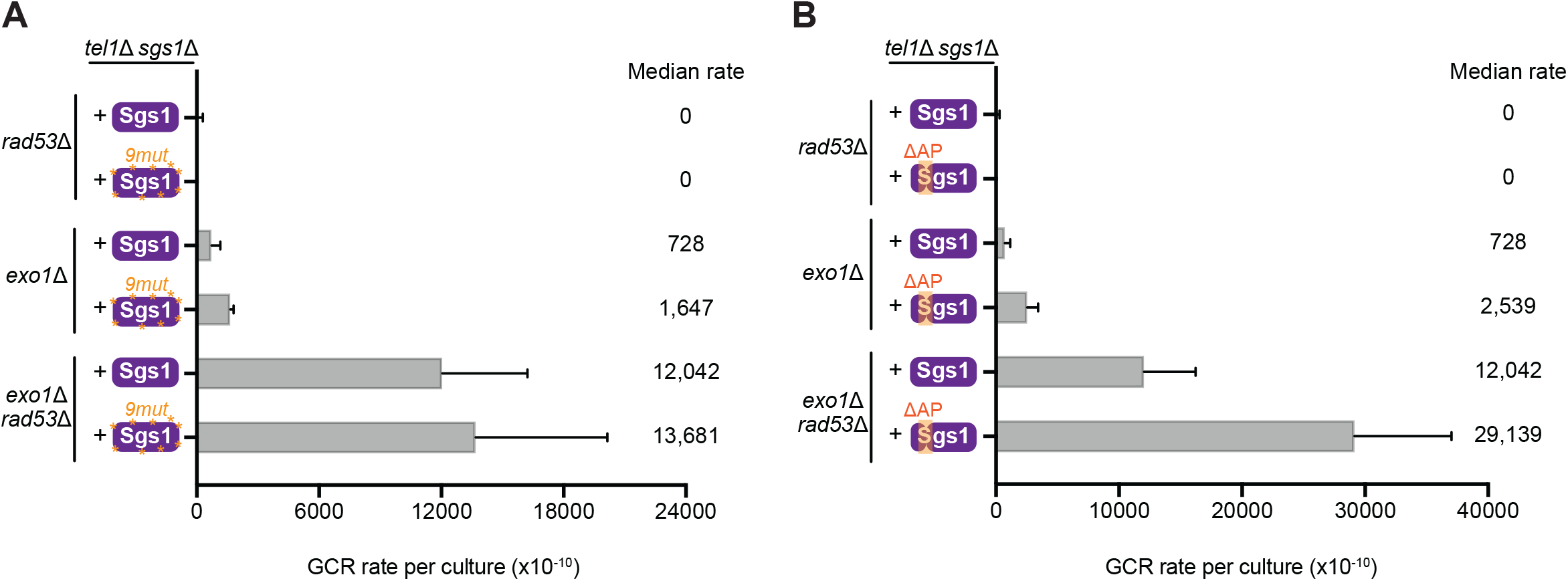
The effect of Sgs1 regulation by Mec1 on GCR suppression in *rad53*Δ cells. (A) Measurement of GCR rates in cells with the indicated genotypes expressing either Sgs1 or Sgs1^9m^^ut^. Bars represent median values and error bars represent standard deviation from 32 independent colonies. (B) Measurement of GCR rates in cells with the indicated genotypes expressing either Sgs1 or Sgs1^APΔ^. Bars represent median values and error bars represent standard deviation from 32 independent colonies.

**Figure S4, related to figure 3.**
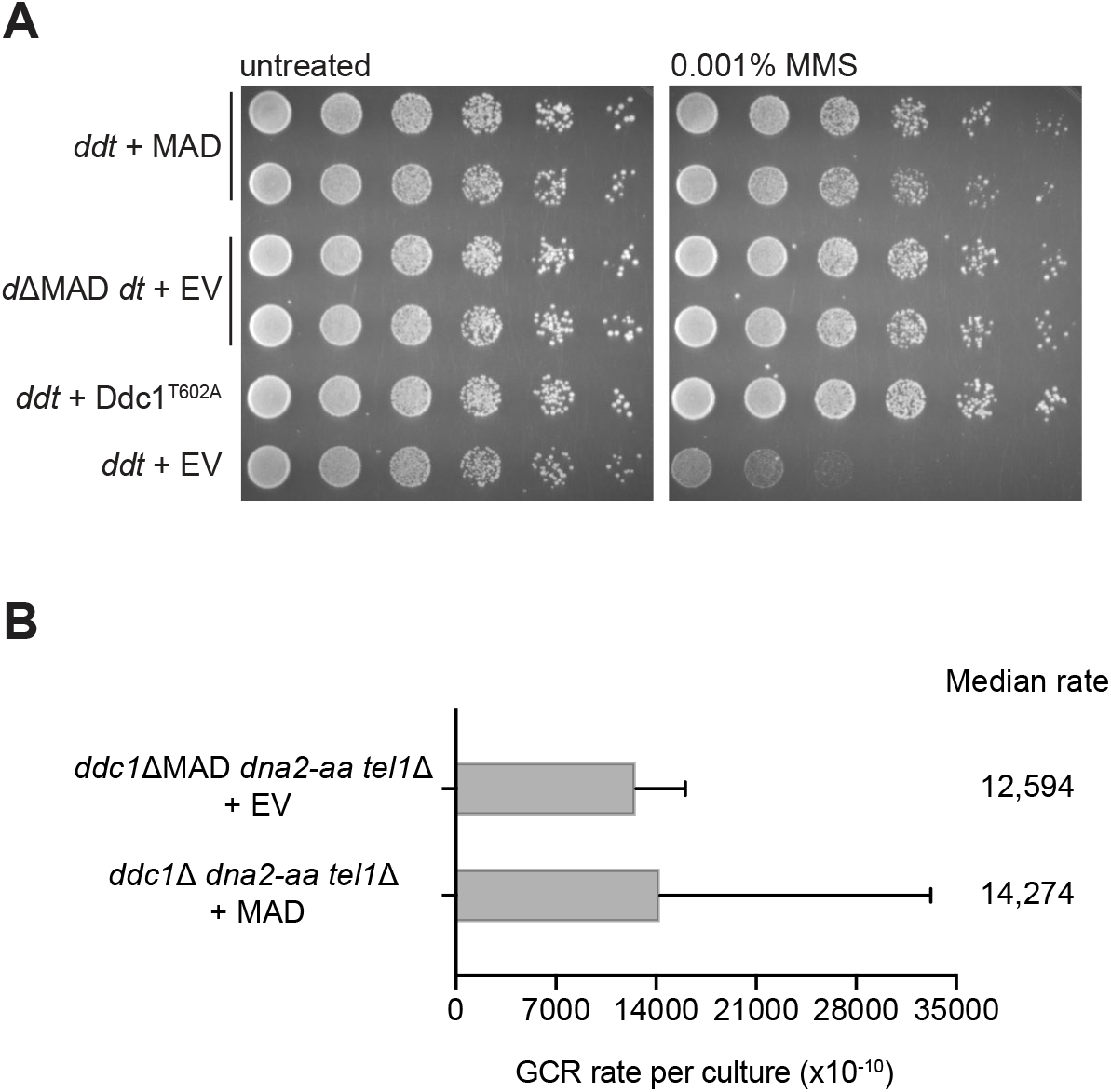
Effects of genomic integration of a Mec1-Activation Domain (MAD) on cell proliferation and GCR rates. (A) Dilution assay of *ddc1*Δ *dna2-aa tel1*Δ (*ddt*) or *ddc1*ΔMAD *dna2-aa tel1*Δ (*d*ΔMAD *dt*) cells expressing either empty vector, MAD or Ddc1^T602A^ in the presence of MMS. 2-fold serial dilutions were used. (B) Measurement of GCR rates in cells with the indicated genotypes expressing either empty vector or MAD. Bars represent median values and error bars represent standard deviation from 32 independent colonies. See Lanz et al., 2018 for more details on the generation of *ddt* cells and effects of MAD expression.

**Figure S5, related to figure 3.**
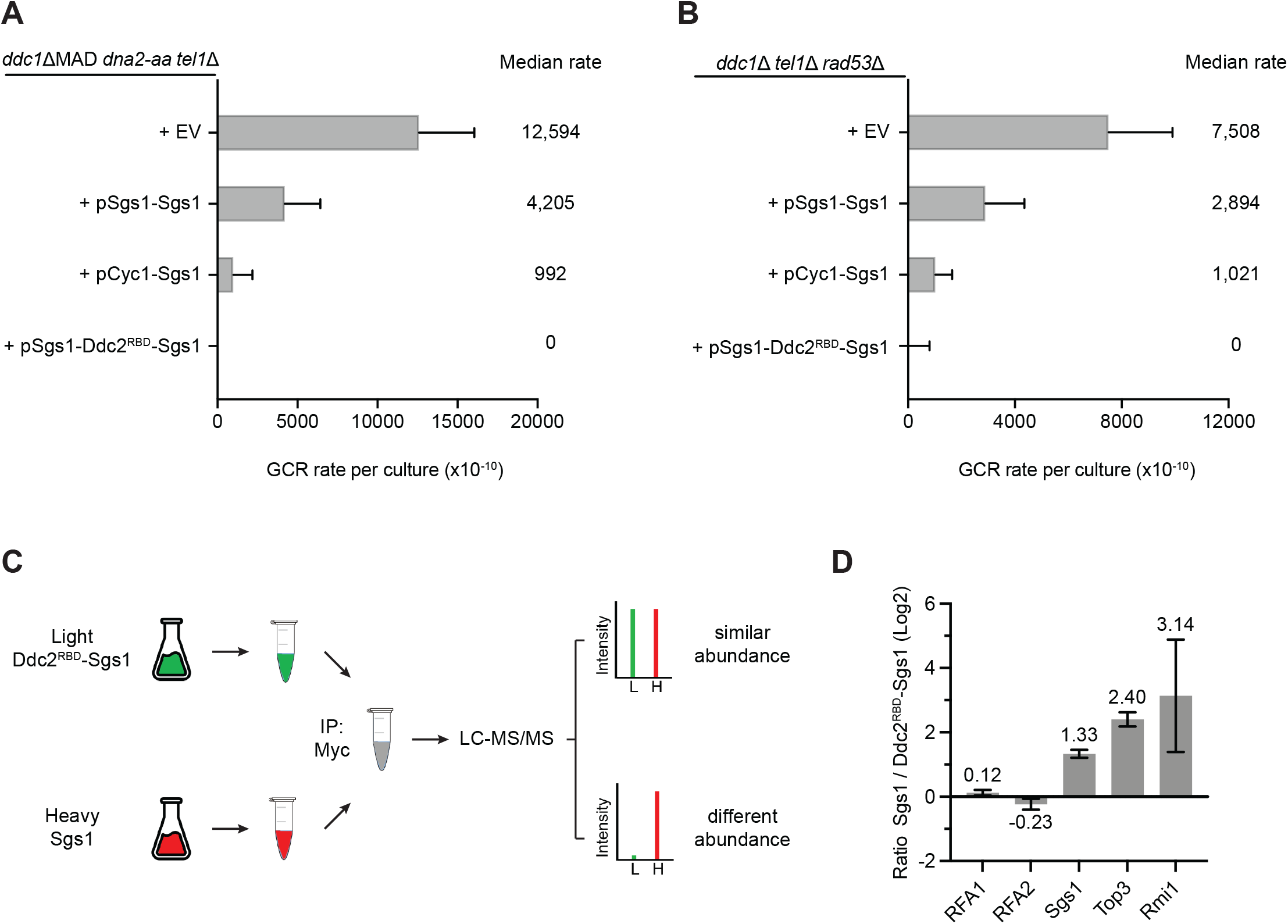
Fusion of RBD to Sgs1 inhibits GCRs independently of abundance changes. (A) Measurement of GCR rates in *ddc1*ΔMAD *dna2-aa tel1*Δ cells expressing empty vector, p*SGS1::SGS1*, p*CYC1::SGS1* and p*SGS1::RBD-SGS1*. Bars represent median values and error bars represent standard deviation from 32 independent colonies. (B) Measurement of GCR rates in *ddc1*Δ *tel1*Δ *rad53*Δ cells expressing empty vector, p*SGS1::SGS1*, p*CYC1::SGS1* and p*SGS1::RBD-SGS1*. Bars represent median values and error bars represent standard deviation from 32 independent colonies. (C) Workflow of the SILAC quantitative mass spectrometry method used to measure the abundance of Sgs1 and RBD-Sgs1. (D) Quantitative mass spectrometry analysis of protein abundance from Sgs1-Myc pull-down experiment. Error bars represent standard deviation of two or more independent peptide-spectrum matches (PSMs) corresponding to the indicated protein. Expression of RBD-Sgs1 is about half of the expression of endogenous Sgs1.

**Figure S6, related to figure 3.**
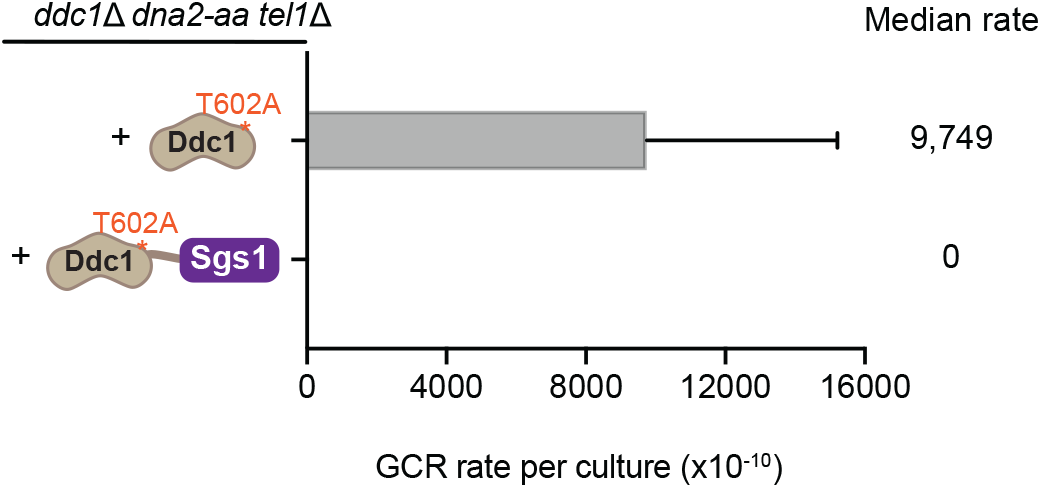
Sgs1 recruitment via fusion with Ddc1 suppresses GCRs in Mec1-deficient cells. Measurement of GCR rates in *ddc1*Δ *dna2-aa tel1*Δ cells expressing either Ddc1^T602A^ or Ddc1^T602A^-Sgs1. Bars represent median values and error bars represent standard deviation from 32 independent colonies.

**Figure S7, related to figure 4.**
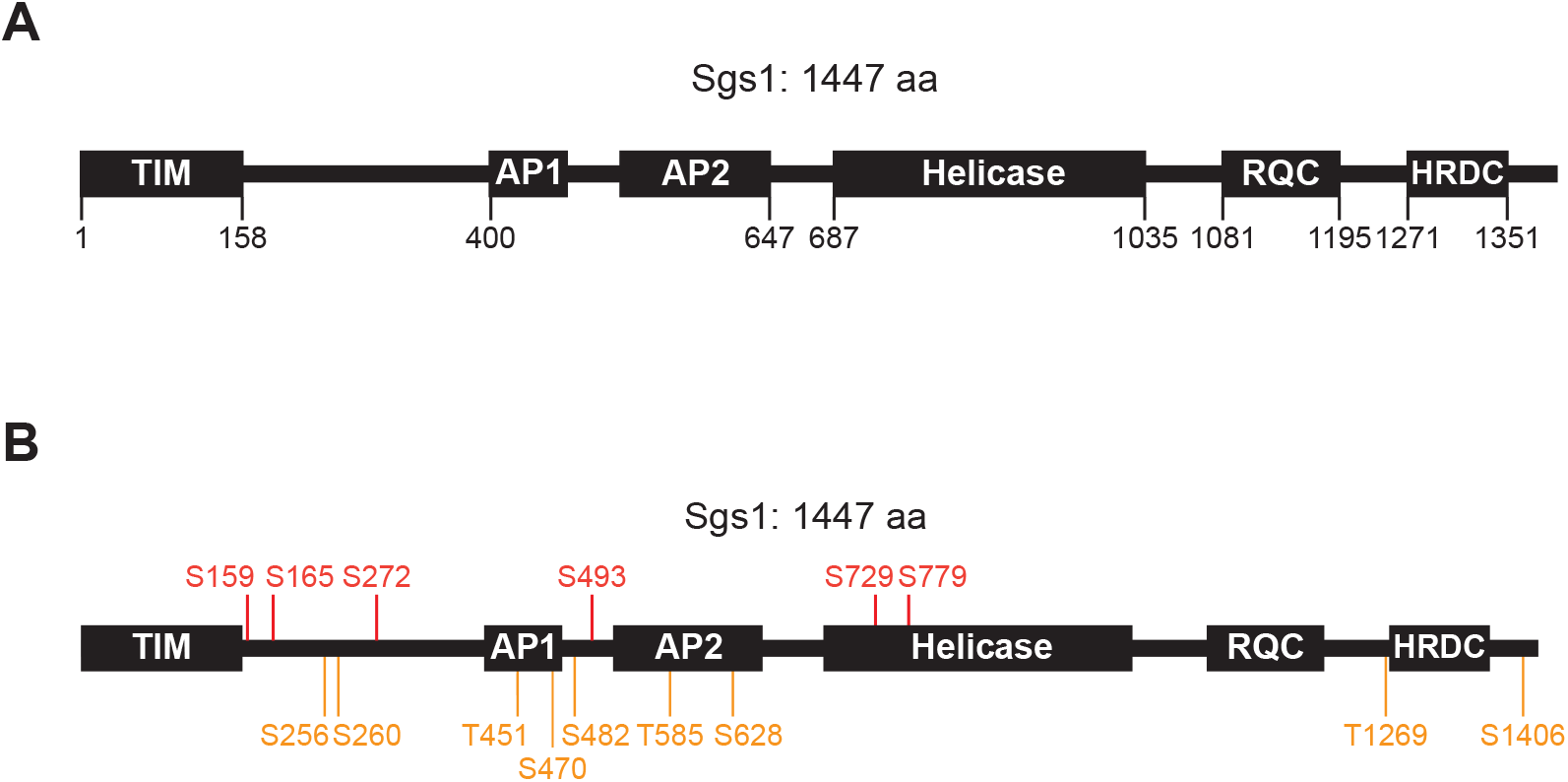
Protein domains and phospho-mutant sites of Sgs1. (A) Schematics depicting Sgs1 domains. (B) Schematics indicating the position of phosphorylation sites mutated in this study. Orange sites represent all SQ/TQ sites (*9mut*) used in this study. Red sites represent four S-P sites (putative CDK motif) and two other non-SQ/TQ sites detected by mass spectrometry, resulting in the *6mut* Mutant.

**Figure S8, related to figure 4.**
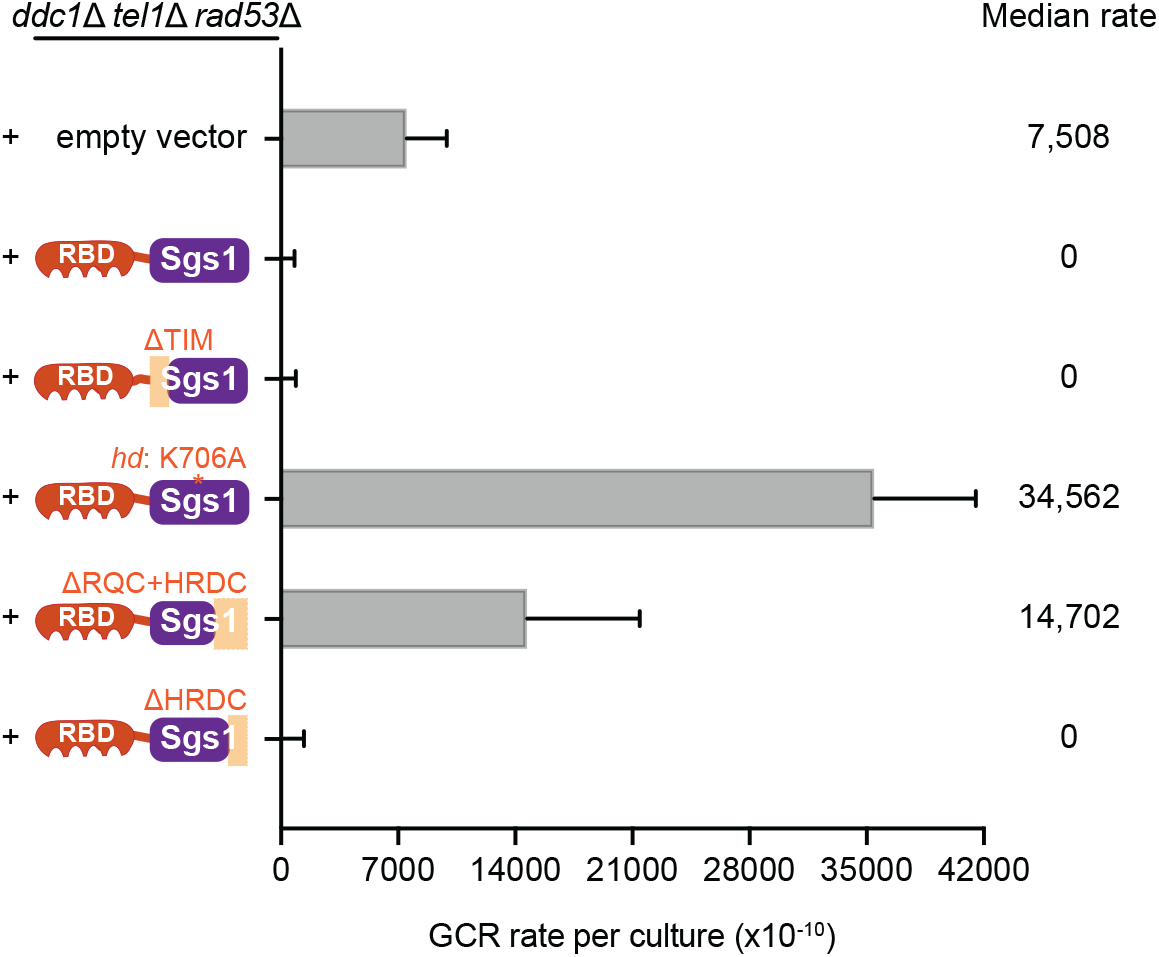
GCR suppression via RBD-Sgs1 requires Sgs1 helicase activity. Measurement of GCR rates in *ddc1*Δ *tel1*Δ *rad53*Δ cells expressing RBD fused to wild-type or truncations of Sgs1. Bars represent median values and error bars represent standard deviation from 32 independent colonies.

**Figure S9, related to figure 5.**
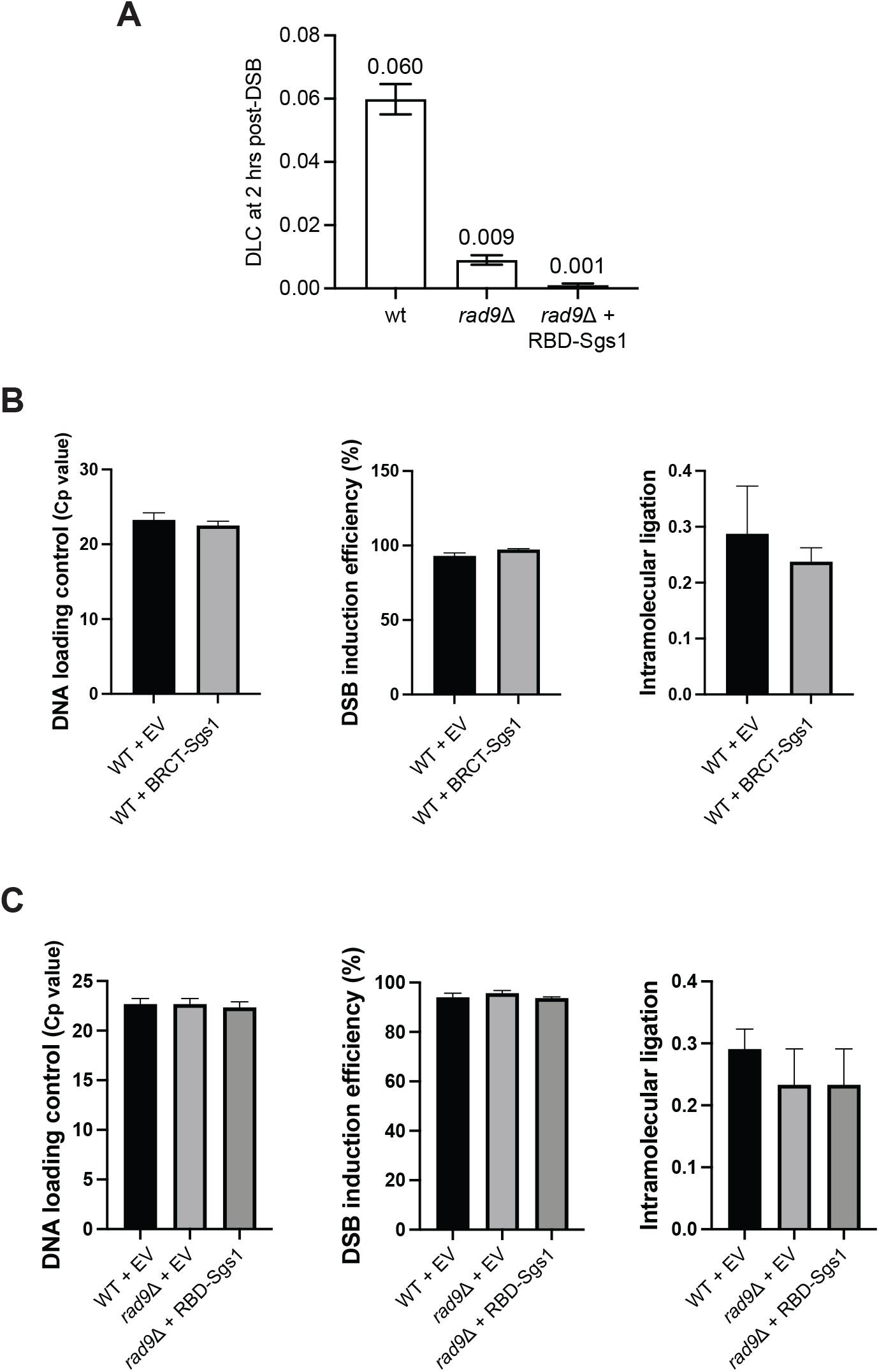
RBD-Sgs1 eliminates D-loop formation. (A) DLC signal in *rad9*Δ cells carrying an empty vector or expressing the RBD-Sgs1 chimera. Error bars represent SEM of two replicate experiments. (B) Control experiments related to figure 5D. (C) Control experiments related to figure S9A.

**Figure S10, related to discussion.**
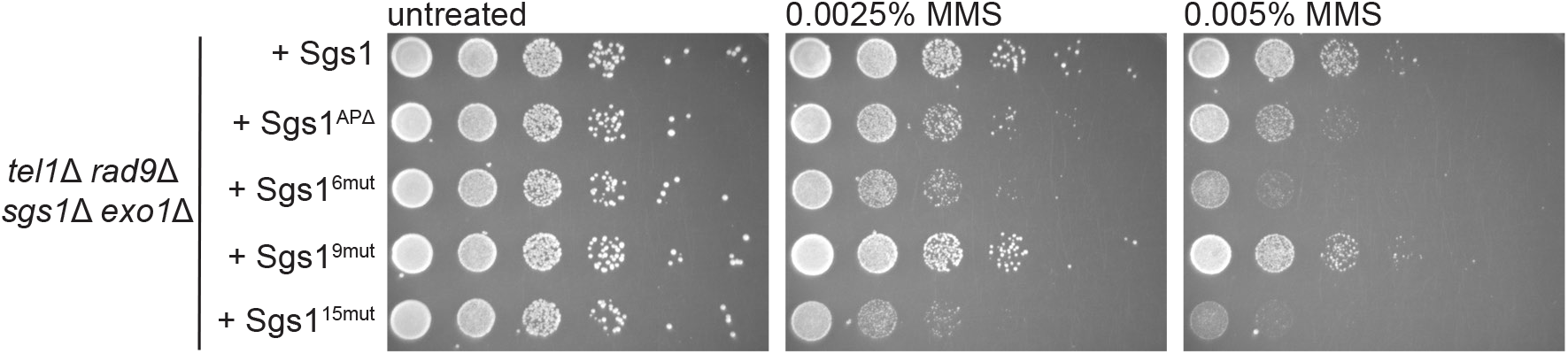
Effects of different Sgs1 mutants on genotoxin response. Dilution assay of *tel1*Δ *rad9*Δ *exo1*Δ *sgs1*Δ cells expressing either wild-type Sgs1 or Sgs1 mutants in the presence of MMS. 10-fold serial dilutions were used.

